# Unlocking a New Path: An Autophagometer that Measures Flux Using a Non-Fluorescent Immunohistochemistry Method

**DOI:** 10.1101/2024.06.26.600741

**Authors:** Shahla Shojaei, Amir Barzegar Behrooz, Marco Cordani, Mahmood Aghaei, Negar Azarpira, Daniel J. Klionsky, Saeid Ghavami

## Abstract

Macroautophagy/autophagy, a crucial cellular process, is typically measured using fluorescence-based techniques, which can be costly, complex, and impractical for clinical settings. In this paper, we introduce a novel, cost-effective, non-fluorescent immunohistochemistry (IHC) method for evaluating autophagy flux. This technique, based on antigen-antibody reactions and chromogenic detection, provides clear, quantifiable results under standard light microscopy, eliminating the need for expensive equipment and specialized reagents. Our method simplifies technical requirements, making it accessible to routine clinical laboratories and research settings with limited resources. By comparing our approach with traditional fluorescence methods, we demonstrate its superior effectiveness, cost-efficiency, and applicability to patient samples. This innovative technique has the potential to significantly advance autophagy research and improve clinical diagnostics, offering a practical and robust tool for studying autophagy mechanisms in diseases such as cancer and neurodegenerative disorders. Our non-fluorescent IHC method represents a significant step forward in evaluating autophagy flux, making it more accessible and reliable, with the promise of enhancing our understanding and treatment of autophagy-related diseases.

## Introduction

Autophagy is an essential cellular mechanism for degrading and recycling components, crucial for maintaining cellular homeostasis and responding to stress [1, 2]. This process is critical in various conditions, including cancer [3, 4], neurodegenerative diseases [5, 6], and infections [7, 8], highlighting its importance to health. Autophagy enables cells to dispose of damaged proteins and organelles, supporting renewal [9, 10]. Autophagy flux is a quantitative measure that assesses the dynamics of this process [11, 12]. Flux tracks the formation of autophagosomes, their fusion with lysosomes, and the degradation of contents. This measure is vital for evaluating the efficiency and regulation of autophagy under different conditions or stimuli. Therefore, monitoring autophagy flux is crucial for understanding the roles of this pathway in biological and disease contexts [13]. However, traditional fluorescence-based techniques have drastic limitations, prompting the development of alternative methods.

Although fluorescence-based methods have been pivotal in advancing our understanding of autophagy, their utility is marred by significant drawbacks. These techniques require sophisticated equipment, specialized reagents, and a high level of technical expertise, escalating the costs and limiting their accessibility to laboratories having substantial resources [14, 15]. Furthermore, these methods prove impractical in clinical settings, where rapid and cost-effective diagnostics are essential. Additionally, their application to fixed tissue samples frequently leads to complications such as tissue autofluorescence, which can mask accurate signals and necessitate extensive sample preparation [16]. This combination of high expense, technical complexity, and practical challenges in clinical environments accentuates the urgent need for more accessible and adaptable methods in autophagy research.

To address these limitations, we propose a novel non-fluorescent immunohistochemistry (IHC) method for evaluating autophagy flux. This approach utilizes antigen-antibody reactions to detect and localize specific antigens, providing invaluable diagnostic, prognostic, and predictive information [17]. We exploit the specificity and robustness of immunohistochemical staining without relying on fluorescence. The specificity is crucial in distinguishing between different autophagic structures, which can overlap in fluorescent studies, while robustness ensures consistent, repeatable results across various sample types and conditions. Using chromogenic detection, this method provides clear and interpretable results that are easily quantifiable under a standard light microscope. The non-fluorescent IHC method is cost-effective and more straightforward than fluorescent approaches, making it accessible to a broader range of laboratories, including those in clinical settings.

This non-fluorescent IHC method offers several significant advantages. First, it reduces costs associated with expensive fluorescent dyes and advanced imaging equipment, eliminating the need for specialized fluorescence microscopes and costly fluorochrome-conjugated antibodies [18]. Second, it simplifies the technical requirements, making it feasible for use in routine clinical laboratories and research settings with limited resources [19]. Third, this method is highly applicable to patient samples, enabling fast and reliable assessment of autophagy flux in various clinical conditions [20]. These features facilitate timely and informed decision-making in clinical diagnostics and research. With this IHC method, we can study autophagy mechanisms in diseases such as cancer and neurodegenerative disorders, where understanding cellular degradation processes is crucial.

This study aims to introduce the non-fluorescent IHC method for evaluating autophagy flux and demonstrate its practical applications across clinical and research settings. By comparing this method with traditional fluorescence-based techniques, we underscore its superior effectiveness, cost-efficiency, and user-friendliness. We anticipate that this innovative approach will transform the assessment of autophagy in patient samples, emerging as a crucial tool for clinical diagnostics and profoundly enhancing our understanding of autophagy-related diseases.

Our innovative non-fluorescent IHC method represents a significant advancement in autophagy research. By offering a cost-effective, accessible, and reliable technique, it holds the potential to transform clinical diagnostics and research, making the evaluation of autophagy flux more practical and widespread.

## Material

### Buffers, Diluent, and Solutions

#### 1. Citrate buffer

Mix 4.5 ml of 0.1 M citric acid (Sigma, 251275-500G) with 20.5 ml of 0.1 M sodium citrate (Sigma, S-4641-500G), and add 225 ml of distilled water (DW) to reach the final volume of 250 ml.

#### 2. Maleic acid buffer

Add 100 mM (116.1 g) maleic acid (Sigma, M0375-500G) and 150 mM (87.5 g) NaCl (Fisher, S271-3) into an Erlenmeyer flask and add water to approximately 800 ml. Adjust the solution to pH 7.5 using 40-70 g of NaOH. After adjusting the pH, increase the volume to 1 liter. **Notes**: i) The maleic acid will only dissolve once the pH reaches 5.5-6. Once it dissolves, add the NaOH pellets very carefully to avoid overshooting the pH. ii) While adjusting the pH, ensure that the temperature is maintained between +15 and +25°C as temperature variations can affect the pH.

#### 3. Blocking stock solution (10X)

Prepare a 10% Blocking Reagent (Roche, 11096176001) in the maleic acid buffer. Dissolve the blocking reagent with heating and shaking while avoiding boiling. Watch carefully to make sure the blocking reagent is completely dissolved. Autoclave and store at 2 to 8°C; if the reagent is stored at -20°C, it can be used for one year.

#### 4. Blocking working solution (1X)

Mix 0.5 ml of blocking stock solution with 1.5 ml of maleic acid buffer, 0.5 ml fetal bovine serum/FBS (Thermofischer, 26140079), 50 μl of 10% Tween-20 (Fisher, BP337-100) and 2.5 ml phosphate-buffered saline (PBS; Milipore-Sigma, 56639). Use a vortex to completely mix.

#### 5. Avidin-Biotin Blocking Solution (Vector, SP-2001)

Ready to use.

### Substrates, Chromogens, and Counterstain Solutions

#### 1. Avidin-Biotin Complex (ABC)

ABC, an avidin-biotin-based enzymatic amplification system, has many biotinylated horseradish peroxidase/HRP molecules cross-linked to avidin, with at least one biotin-binding site able to bind to the secondary antibody and amplify the signal. Prepare ABC solution by adding 1 drop from bottle “A” (Vector, PL-6100) and 1 drop from bottle “B” to 2.5 ml of PBS. Leave the solution for 30 min at room temperature to be activated. **Note:** This solution is stable for 1 week at 4^°^C.

#### 2. Peroxidase chromogen kit, diaminobenzidine (DAB)

Prepare DAB substrate by adding 125 μl of 1% DAB (ThermoFisher, 34002) stock solution and 50 μl of 30% H_2_O_2_ to 2.5 ml PBS.. **Note:** The substrate should be prepared fresh just before use. **See Notes and Tips 1**.

#### 3. Counterstain

Add 10 drops of Mayer hematoxylin (Vector, H-3404) to 1.25 ml of PBS.

#### 4. Bluing reagent for counterstaining

Prepare 2% sodium bicarbonate (Fisher Scientific, BP328-1) in DW.

### Antibodies

#### 1. Anti-SQSTM1/p62 (D5L7G) primary antibody

Prepare 1:50 dilution of SQSTM1/p62 (sequestosome 1) antibody (Cell Signaling Technology, 88588) in blocking working solution. **Note:** Primary antibody concentration may vary depending on the specific system being used. To optimize primary antibody concentration in your system, perform a serial dilution around the recommended starting concentration. **See Notes and Tips 2**.

#### 2. Anti-MAP1LC3B/LC3B-primary antibody

Prepare 1/50 dilution of MAP1LC3B/LC3B (microtubule-associated protein 1 light chain 3 beta)-Specific Antibody (ProteinTech, 18725-1-AP) in blocking working solution. **Note:** The primary antibody concentration may vary depending on the specific system being used. To optimize the primary antibody concentration in your system, perform a serial dilution around the recommended starting concentration.

#### 3. Biotinylated anti-mouse secondary antibody

IHC Select Secondary Goat Anti-Mouse IgG Antibody, prediluted, biotinylated (Sigma-Aldrich, 21538-M).

#### 4. Biotinylated anti-rabbit secondary antibody

IHC Select Secondary Goat Anti-Rabbit IgG Antibody, prediluted, biotinylated (Sigma-Aldrich, 21537).

### Miscellaneous

#### 1. Silanized slides

To prevent tissue detachment, use silanized (positively charged) slides (Sigma, S4651).

#### 2. Mounting medium

To form a semi-permanent seal for prolonged slide storage at 4°C use Flourmount-G (Invitrogen, 00-4958-02).

#### 3. Xylene, histological grade

For dehydration and rehydration steps, use ready-to-use xylene (Sigma Aldrich, 534056).

#### 4. Ethanol

For dehydration and rehydration steps, use food-grade ethanol with high purity (UN 1170; Greenfield Global, PO25EAAN).

### Method

Unless otherwise specified, perform all steps at room temperature.

#### Day 1

##### 1. De-paraffinize and rehydrate slides

Carry out the following exchange at room temperature, manually in Coplin jars (Millipore-Sigma, S5641), or using an automated embedding system. **See Notes and Tips 3**.

- Warm the slides at 60^°^C for 30 min in the Coplin jar.
- Place the slides in xylene for 5 min (3X), being sure to cover the samples in this and all subsequent steps.
- Place the slides in 100% ethanol for 1 min (2X).
- Place the slides in 95% ethanol for 1 min.
- Place the slides in 70% ethanol for 1 min.
- Wash with running tap water for 1 min.
- Place the slides in PBS for 1 min (3X).

##### 2. Heat-induced antigen retrieval

- Pre-warm citrate buffer in a polypropylene Coplin jar. **See Notes and Tips 4**.
- Place the slides in the Coplin jar.
- Place the Coplin jar in a boiling water bath for 30 min.
- Remove the Coplin jar from the boiling water.
- Allow to cool slowly at room temperature for 20 min.

##### 3. Blocking of endogenous proteins

Blocking steps are crucial in IHC to prevent excessive background staining in images. **See Notes and Tips 5**.

- Remove slides from the citrate buffer one by one and dip quickly in double-distilled H_2_O.
- Gently blot on a paper tissue to remove liquid.
- Circle the tissue sections with an ImmEdge pen (Vector, H-4000).
- Immediately cover the tissue sections with PBS. **Note**: Do not allow the sections to dry.
- Apply PBS to the sections for 5 min to wash (3X).
- Apply the blocking solution to the sections for 30 min.
- Apply PBS to the sections for 5 min to wash (3X). **Note:** If you are using an antibody produced in the same species as your tissue (e.g., a mouse antibody on mouse tissue), blocking endogenous IgG is necessary. For example, use goat anti-mouse IgG Fab fragment (Jackson ImmunoResearch, 115-007-003) diluted 1:10 in PBS for a minimum of 1 h. It is best to incubate overnight at 4°C. Then, wash sections in PBS for 5 min (3X). If this is not an issue, proceed to the next step.
- Apply freshly prepared 3% H_2_O_2_ (in PBS) for 10 min. This step is necessary to eliminate endogenous hydrogen peroxidases. **Note:** The incubation time may have to be increased to 15 or 20 min if the tissue has a lot of blood.
- Apply PBS to the sections for 5 min to wash (3X).
- Apply avidin blocking solution for 15 min. **Note**: This solution works just as well when diluted in an equal volume of PBS.
- Apply PBS to the sections for 5 min to wash (3X).
- Apply biotin-blocking solution for 15 min. **Note**: This solution works just as well when diluted in an equal volume of PBS.
- Apply PBS to the sections for 5 min to wash (3X).

##### 4. Immunostaining

- Arrange sections in a moist chamber to avoid drying of tissues.
- Apply primary antibody and incubate overnight at 4°C. **Note:** Leave one section without primary antibody, and cover only with blocking solution to be used as a negative control. **See Notes and Tips 7**.

#### Day 2

- Apply PBS to the sections for 5 min to wash (3X).
- Apply biotinylated secondary antibody (corresponding to the primary antibody) to all sections (including the negative control) for 30 min.
- Apply PBS to the sections for 5 min to wash (3X).
- Apply activated “ABC” solution for 30 min.
- Apply PBS to the sections for 5 min to wash (3X).
- Apply freshly prepared DAB substrate to the sections for up to 2 min. **See Notes and Tips 8**.
- Stop reaction by immersing slide in double-distilled H_2_O in a Coplin jar.
- Wash slides with H_2_O in a Coplin jar (3X).
- Counterstain with Mayer hematoxylin for 1-4 min. **See Notes and Tips 9**.
- Wash slides with H_2_O in a Coplin jar (3X),
- Immerse in 2% sodium bicarbonate solution for 20 s.
- Wash slides with H_2_O in a Coplin jar (3X).

##### 5. Dehydration and mounting

Dehydration is done by reversing the steps of hydration. Carry out the following exchange at room temperature, manually in Coplin jars, or using an automated embedding system. **See Notes and Tips 10**.

- Place the slides in 70% ethanol for 1 min.
- Place the slides in 95% ethanol for 1 min.
- Place the slides in 100% ethanol for 1 min (2X).
- Place the slides in xylene for 5 min (3X).
- Remove slides one by one.
- Blot the edge of the slide to a paper tissue to remove the extra liquid.
- Add one drop of mounting medium on the top of the tissue section.
- Mount a coverslip on the mounting medium.
- Gently press the coverslip with the tip of a pencil to remove any possible bubbles.
- Lightly clean the extra mounting agent surrounding the coverslip using the edges of a paper tissue.
- Leave slides at room temperature overnight to ensure they are completely dry.

#### Day 3

##### Imaging

To perform microscopy imaging for DAB IHC, tissue sections are prepared and stained with primary and secondary antibodies, followed by a DAB substrate, then mount the stained sections on slides and cover them with coverslips as described above. Use a bright-field microscope equipped with a camera. Adjust the light and focus to visualize the brown DAB precipitate. Capture images at multiple magnifications (e.g., 10x, 20x, 40x). Ensure consistent imaging settings for all samples. Save and label images appropriately for analysis. This protocol provides clear visualization of DAB-stained tissue sections, facilitating accurate evaluation and documentation.

To ensure unbiased and accurate results, scoring DAB IHC for LC3 and SQSTM1 puncta involves a meticulous evaluation by three blinded pathologists. Each pathologist independently examines stained tissue sections under a microscope, focusing on the presence and intensity of the brown DAB residue, which indicates the presence of the target antigens LC3 and SQSTM1. The scoring process includes assessing the puncta staining intensity and the percentage of positively stained cells. The puncta staining intensity for LC3 and SQSTM1 is graded on a scale from 0 to 3, where 0 indicates no puncta staining (null), 1 indicates weak puncta staining (low), 2 indicates moderate puncta staining (medium), and 3 indicates strong puncta staining (high). The percentage of positively stained cells is categorized into ranges (e.g., 0-25%, 26-50%, 51-75%, and 76-100%). Each pathologist assigns scores independently, unaware of the others’ assessments or identifying information about the samples. This blinded approach minimizes bias and enhances the reliability of the results. After scoring, the results from the three pathologists are compared and averaged to obtain a final score for each sample. This rigorous evaluation method ensures that the interpretation of LC3 and SQSTM1 puncta via DAB-based IHC staining is accurate and reproducible, providing reliable data for research and diagnostic purposes. This process is critical for studies investigating autophagy, as LC3 and SQSTM1 are key markers of this cellular process.

#### Day 5

##### Autophagy Flux Evaluation, Overview

Evaluating autophagy flux involves assessing the expression levels and patterns of cytosolic LC3 and SQSTM1 puncta, which are crucial markers of autophagy activity. Autophagy flux refers to the dynamic autophagosome formation, maturation, and degradation process. Note that there is an ongoing balance between LC3 and SQSTM1 synthesis and degradation. During autophagy induction, LC3 levels in particular increase. LC3 is present on both sides of the phagophore, the initial sequestering compartment, and the autophagosome. The LC3 on the outer surface of the autophagosome is removed and recycled, whereas the population on the inner surface is exposed to the lysosome lumen and degraded, lowering the LC3 level. SQSTM1 is bound to LC3 on the inner surface of the phagophore and autophagosome and hence is also degraded in the lysosome. Therefore, an increase in LC3 can reflect i) autophagy induction or ii) a block in autophagosome-lysosome fusion or degradation within the lysosome. Accordingly, measuring LC3 alone is not sufficient to assess autophagic flux. In general, an accumulation of LC3 along with SQSTM1 would indicate a block in flux. In contrast, an increase in LC3 accompanied by a decrease in SQSTM1 could indicate high autophagy activity (a large increase in LC3 due to autophagy induction along with turnover of both proteins). Thus, the balance between the levels of LC3 and SQSTM1 provides substantial information about flux.

#### Specific Interpretation of Puncta Formation

##### 1. High LC3 and SQSTM1 Puncta

Indicates **inhibition of autophagy flux**. Both markers are elevated because autophagy is induced and autophagosomes are formed but not degraded (**Fig. 1A**).

**Figure 1.**
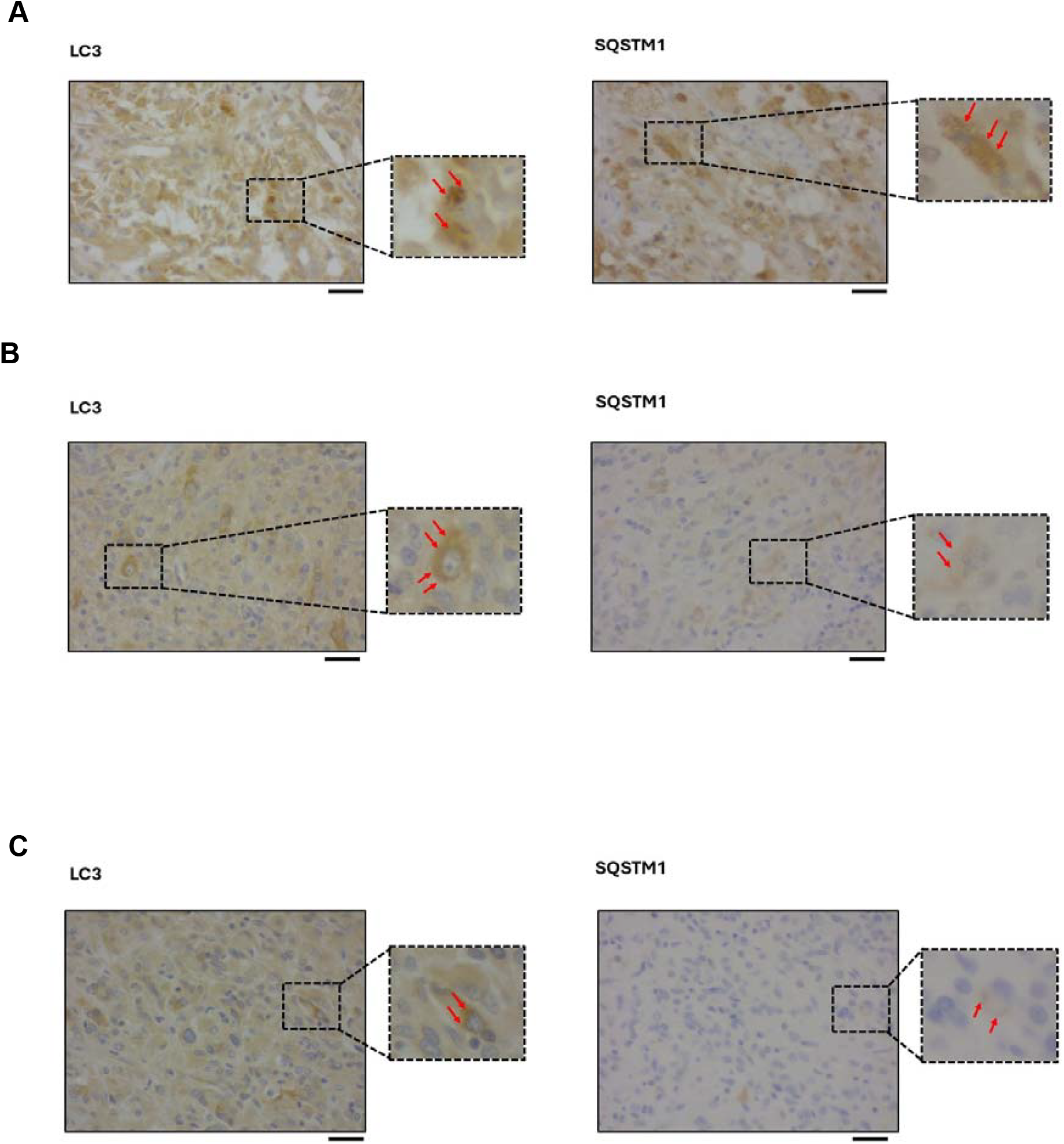
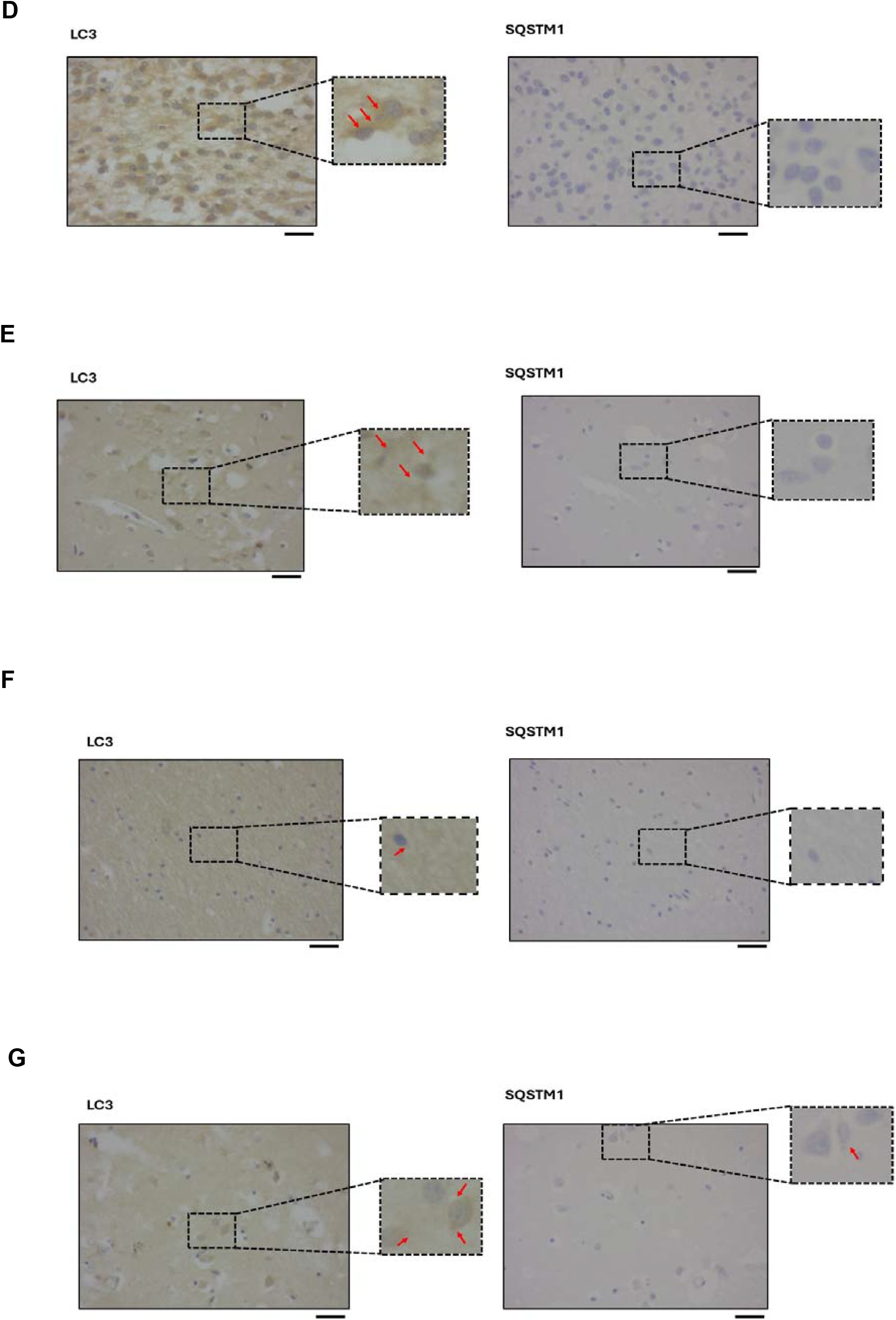
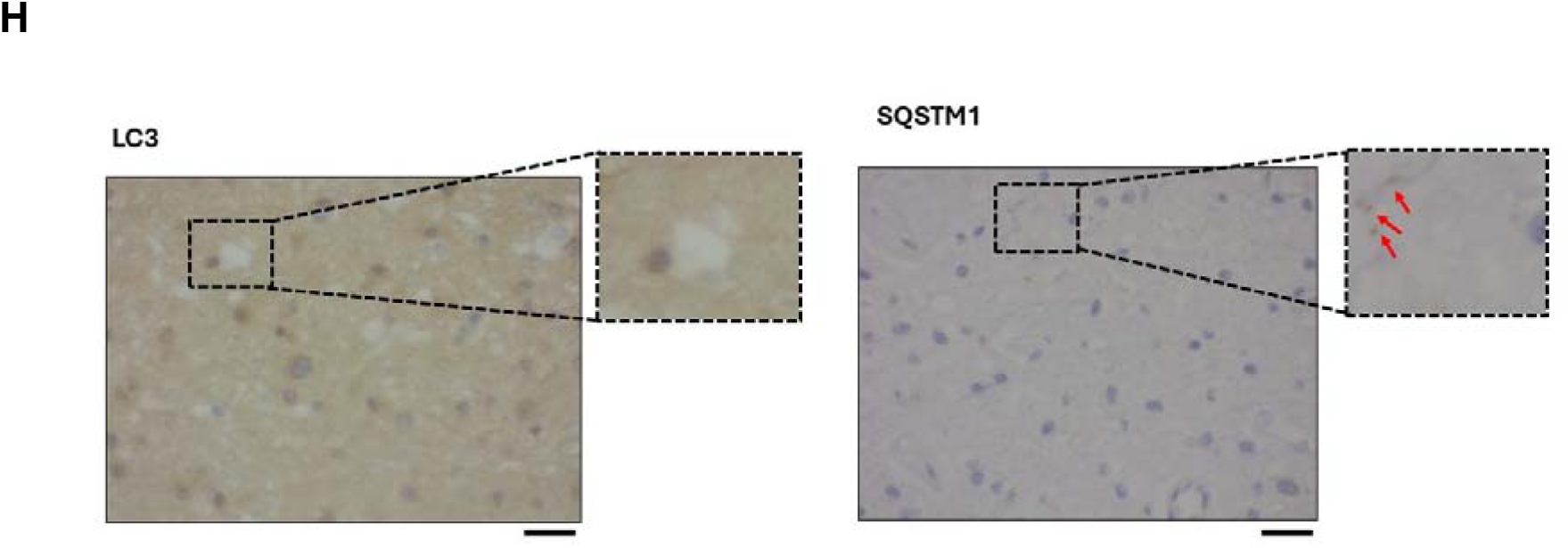
IHC analysis of autophagy flux in human cerebrum tissue. **(A)** Autophagy flux inhibited. Immunohistochemical staining shows high expression levels of both LC3 and SQSTM1. **(B)** Autophagy flux is moderately inhibited. IHC staining shows high expression levels of LC3 and medium expression of SQSTM1. **(C)** Very Active Autophagosome Formation. IHC staining shows high expression levels of LC3 and low expression of SQSTM1. **(D)** Very Active Autophagosome Formation. IHC staining shows high expression levels of LC3 and null expression of SQSTM1. **(E)** Very Active Autophagic Flux. IHC staining shows medium expression levels of LC3 and null expression of SQSTM1. **(F)** Very Active Autophagic Flux. IHC staining shows low expression levels of LC3 and null expression of SQSTM1. **(G)** Very Active Autophagic Flux. IHC staining shows low expression levels of both LC3 and SQSTM1. **(H)** Very Active Autophagic Flux. IHC staining shows null expression levels of LC3 and low expression of SQSTM1.

##### 2. High LC3 and Medium SQSTM1 Puncta

Suggests **moderate autophagy flux inhibition**. This scenario shows some autophagic activity but incomplete degradation of autophagosomes (**Fig. 1B**).

##### 3. High LC3 and Low/Null SQSTM1 Puncta

Reflects **very active autophagosome formation**. High LC3 indicates numerous autophagosomes, while low SQSTM1 suggests efficient degradation:

∘ (I) Autophagosome formation is highly active with subsequent degradation (**Fig. 1C**).
∘ (II) Autophagosome formation is highly active without accumulation of SQSTM1 (**Fig. 1D**).

##### 4. Medium LC3 and Null SQSTM1 Puncta

Denotes **very active flux (I)**. This indicates a balanced and efficient autophagic process where autophagosomes are consistently formed and degraded (**Fig. 1E**).

##### 5. Low LC3 and Null SQSTM1 Puncta

Signifies **very active flux (II)**. Here, the autophagic process is so efficient that few autophagosomes accumulate, and SQSTM1 is rapidly degraded (**Fig. 1F**). **See Notes and Tips 6**.

##### 6. Low LC3 and Low SQSTM1 Puncta

Corresponds to **very active flux (III)**, indicating effective autophagosome formation and turnover (**Fig. 1G**).

##### 7. Null LC3 and Low SQSTM1 Puncta

Suggests **very active flux (IV)**. This scenario represents a state where autophagosomes are scarce due to their rapid turnover, and SQSTM1 levels remain low (**Fig. 1H**).

By carefully monitoring these patterns, researchers can infer the efficiency of the autophagic process, which is vital for understanding cell health and responses to various treatments or stressors. For a summary, please see **Table 1**.

**Table 1.** Autophagy flux indicators and interpretation.

| IHC staining score of autophagy flux indicators |  | IHC staining interpretation |
| --- | --- | --- |
| Cytosolic LC3 puncta | Cytosolic SQSTM1 puncta |  |
| High<sup>a</sup> | High | Autophagy flux inhibited |
| High | Medium | Autophagy flux moderately inhibited |
| High | Low | Very active autophagosome formation (I) |
| High | Null | Very active autophagosome formation (II) |
| Medium | Null | Very active flux (I) |
| Low<sup>b</sup> | Null | Very active flux (II) |
| Low | Low | Very active flux (III) |
| Null | Low | Very active flux (IV) |

<sup>a</sup>High: 3; medium: 2; low: 1; null: 0
<sup>b</sup>See Notes and Tips 6.

### Notes and Tips

Here are ten important notes and tips for achieving optimal results with DAB IHC to measure autophagy flux:

#### 1. Use Fresh Reagents

Ensure all reagents, especially DAB substrate, hydrogen peroxide, and antibodies, are fresh. DAB is light-sensitive and can degrade, leading to reduced staining efficiency.

#### 2. Optimize Antibody Concentrations

Finding the right dilution of primary and secondary antibodies is crucial for specific and strong staining. Use titration to determine the optimal antibody concentration for your specific tissue and antigen.

#### 3. Proper Tissue Fixation

Over-fixation or under-fixation can adversely affect antigen availability. Standardize fixation time and conditions. Typically, 10% formalin for 24 h is effective for most tissues.

#### 4. Antigen Retrieval

Perform appropriate antigen retrieval methods, either heat-induced (using a microwave, pressure cooker, or steamer) or enzymatic, to unmask epitopes. Use a retrieval buffer like citrate or ethylenediaminetetraacetic acid/EDTA at the correct pH.

#### 5. Control Background Staining

To minimize non-specific staining, block endogenous peroxidase activity with hydrogen peroxide (usually 0.3% in methanol). Use normal serum or bovine serum albumin/BSA to block non-specific binding sites.

#### 6. Use Appropriate Controls

Always include positive and negative controls. These include a tissue known to express the antigen (positive control) and a sample in which the primary antibody is omitted or replaced with an irrelevant antibody (negative control). Low levels of LC3 may indicate efficient turnover of the protein, or a low level of autophagy induction. The extent of LC3 upregulation also varies depending on the tissue or cells being evaluated. Thus, in situations where there are low levels of both LC3 and SQSTM1 we recommend caution in interpretation of the results unless additional methods of evaluation are performed. For researchers using cell culture appropriate controls will include the addition of lysosomal protease inhibitors such as E64d and leupeptin with and without an MTOR inhibitor such as torin 1, which will induce autophagy. The combination of MTOR and protease inhibitors will indicate the highest levels of LC3 and SQSTM1 that can be obtained in the particular cell lines being used. See reference [21] [X] for additional information.

#### 7. Optimized Incubation Times and Temperatures

Incubation times and temperatures for primary and secondary antibodies should be optimized. Overnight incubation at 4°C for the primary antibody often enhances specificity.

#### 8. Monitor DAB Development Time

The development time for DAB is critical. Overdevelopment can lead to non-specific staining and high background. Monitor under a microscope and stop the reaction by washing with water as soon as the desired intensity is achieved.

#### 9. Counterstaining

Use an appropriate counterstain (e.g., hematoxylin) to provide contrast and enable better visualization of tissue morphology alongside DAB staining. Optimize the counterstaining time to avoid obscuring DAB signals.

#### 10. Proper Mounting

Use aqueous or permanent mounting media compatible with DAB. Ensure that slides are properly dried and coverslipped to prevent fading and preserve the staining.

These tips can significantly improve the reliability and quality of DAB-based IHC staining, ensuring clear and interpretable results.

## Funding

M.C. is supported by grant RYC2021-031003I funded by MICIU/AEI/https://doi.org/10.13039/501100011033 and, by European Union NextGenerationEU/PRTR; DJK is funded by NIH grant GM131919. SG was supported by CCMF Operating grant (763117252).

## Conflict of interest

The authors declare no conflict of interest.

## Author contributions

Shahla Shojaei and Mahmoud Aghaei: Methodoogy, preparation of draft’ Amir Barzegar Behrooz, and Marco Cordani: Figure preparation, preparation of draft; Negar Azarpira: pathological evaluation and scoring (leader of pathology team); Daniel J. Klionsky: evaluation of the results and final edit; Saeid Ghavami: innovation of method, designing the method, resources, supervision, finalizing the results.

## Data Accessibility

All raw data are accessible upon request.

Ethic: All tissue that were used are commercial TMA from US Biomax.

### Abbreviations

ABC: avidin-biotin complex
DAB: diaminobenzidine
DW: distilled water
IHC: immunohistochemistry
LC3: microtubule-associated protein 1 light chain 3 beta
NaOH: aodium hydroxide
PBS: phosphate-buffered saline
PMSF: phenylmethylsulfonyl fluoride
SDS: sodium dodecyl sulfate
SQSTM1: sequestosome 1.

## Notes

### Competing Interest Statement

The authors have declared no competing interest.

